# Small molecules with antibiofilm, antivirulence and antibiotic synergy activities against *Pseudomonas aeruginosa*

**DOI:** 10.1101/067074

**Authors:** Erik van Tilburg Bernardes, Laetitia Charron-Mazenod, David Reading, Shauna L. Reckseidler-Zenteno, Shawn Lewenza

## Abstract

Biofilm formation is a universal bacterial strategy for long-term survival in nature and during infections. Biofilms are dense microbial communities enmeshed within a polymeric extracellular matrix that protects bacteria from antibiotic exposure and the immune system and thus contribute to chronic infections. *Pseudomonas aeruginosa* is an archetypal biofilm-forming organism that utilizes a biofilm growth strategy to cause chronic lung infections in Cystic Fibrosis (CF) patients. The extracellular matrix of *P. aeruginosa* biofilms is comprised mainly of exopolysaccharides (EPS) and DNA. Both mucoid and non-mucoid isolates of *P. aeruginosa* produces the Pel and Psl EPS, each of which have important roles in antibiotic resistance, biofilm formation and immune evasion. Given the central importance of the Pel and Psl EPS in biofilm structure, they are attractive targets for novel anti-infective compounds. In this study we used a high throughput gene expression screen to identify compounds that repress expression of *pel* and *psl* genes as measured by transcriptional *lux* fusions. Testing of the *pel/psl* repressors demonstrated an antibiofilm activity against microplate and flow chamber biofilms formed by wild type and hyperbiofilm forming strains. To determine the potential role of EPS in virulence, mutants in *pel/psl* were shown to have reduced virulence in the feeding behavior and slow killing virulence assays in *Caenorhabditis elegans*. The antibiofilm molecules also reduced *P. aeruginosa* PAO1 virulence in the nematode slow killing model. Importantly, the combination of antibiotics and antibiofilm compounds were synergistic in killing *P. aeruginosa* biofilms. These small molecules represent a novel anti-infective strategy for the possible treatment of chronic *P. aeruginosa* infections.

**Author summary:** Bacteria use the strategy of growing as a biofilm to promote long-term survival and therefore to cause chronic infections. One of the best examples is *Pseudomonas aeruginosa* and the chronic lung infections in individuals with Cystic Fibrosis (CF). Biofilms are generally a dense community of bacteria enmeshed in an extracellular matrix that protects bacteria from numerous environmental stresses, including antibiotics and the immune system. In this study we developed an approach to identify *P. aeruginosa* biofilm inhibitors by repressing the production of the matrix exopolysaccharide (EPS) polymers. Bacteria treated with compounds and then fed to the nematode also had showed reduced virulence by promoting nematode survival. To tackle the problem of biofilm tolerance of antibiotics, the compounds identified here also had the beneficial property of increasing the biofilm sensitivity to different classes of antibiotics. The compounds disarm bacteria but they do not kill or limit growth like antibiotics. We provide further support that disarming *P. aeruginosa* may be a critical anti-infective strategy that limits the development of antibiotic resistance, and provides a new way for treating chronic infections.

## Introduction

Biofilm formation is a universal virulence strategy adopted by bacteria to survive in hostile environments [1, 2]. The Gram-negative opportunistic pathogen *Pseudomonas aeruginosa* is a remarkable biofilm-forming species that commonly establishes chronic infections in the lungs of patients with the genetic disease Cystic Fibrosis (CF) [2, 3]. Growth as a biofilm promotes multidrug resistance to antibiotic interventions and evasion of immune clearance [1, 4, 5]. Biofilm formation is a conserved process of attachment, maturation and dispersion, where sessile, bacterial aggregates are held together by a protective polymeric extracellular matrix comprised mainly of exopolysaccharides (EPS) and extracellular DNA [1, 4, 6–8]. *P. aeruginosa* strains produce three different EPS molecules; alginate, Pel and Psl [9]. Pel and Psl are the major EPS produced in the early CF colonizing, non-mucoid isolates [10, 11], and also contribute to biofilm formation in mucoid CF isolates, which overproduce alginate and emerge as the infection progresses [8, 12].

Both Pel and Psl have diverse roles in biofilm formation, antibiotic resistance, immune evasion, and whose overproduction leads to hyperaggregative small colony variants (SCVs) [1, 13]. Pel is a positively charged EPS, formed by partially acetylated galactosamine and glucosamine residues, with both cell-associated and secreted forms [5]. Psl is a neutrally charged EPS, comprised of repeating pentamers of D-mannose, D-glucose and L-rhamnose, which can be also found as part of the bacterial capsule and secreted to form the biofilm matrix [6, 14]. Both Pel and Psl are able to initiate biofilm formation [1, 15]. Pel functions as an adhesin that is critical for initial cell-cell and cell-surface interactions and the formation of pellicles in the air-liquid interface [1, 15]. Pel also has a structural role in cross-linking eDNA, establishing the scaffold of the biofilm [5]. In the *Drosophila melanogaster* oral feeding model, Pel is highly expressed and required for biofilm formation in the fruit fly crop [16]. Mutation in the *pel* operon results in rapid escape from the gastrointestinal tract and faster killing of *D. melanogaster*, highlighting the function of EPS to limit dissemination in chronic fruit fly infections [16]. Psl arranges in fiber-like structures that are also crucial for cell-surface interactions, matrix development and biofilm architecture [1, 6, 7]. Both Pel and Psl are also involved in antimicrobial resistance, where Pel is crucial for increased biofilm resistance to aminoglycosides [15] and Psl contributes short-term tolerance to polymyxins, aminoglycosides and fluoroquinolone antibiotics [17]. Further, Psl has also been shown to reduce recognition by the innate immune system, blocking complement deposition on the bacterial surface and reducing phagocytosis, release of reactive oxygen species (ROS) and cell killing by neutrophils [18].

Biofilms are intimately related to antibiotic tolerance and persistent infections [15, 19], therefore, there is an urgent need for the identification of new approaches that target and inhibit the biofilm mode of growth for the prevention or treatment of chronic bacterial infections. In order to identify new molecules effective against biofilms, high-throughput screening (HTS) approaches have been employed to screen large numbers of compounds that reduce biofilm formation and/or detach pre-formed biofilms in many species of bacteria [20–24].

Given the importance of Pel and Psl in *P. aeruginosa* biofilm formation, they are attractive targets for antibiofilm drug development. In this study we used a HTS gene expression approach to screen a 31,096-member small-molecule drug library for compounds that repress *pel* and *psl* gene expression. Consistent with our hypothesis, the *pel/psl* repressor compounds inhibited EPS secretion and also had significant antibiofilm activity. Further testing of these compounds revealed their antivirulence activity in *Caenorhabditis elegans* infection model, and their synergy with conventional antibiotics. The anti-infective compounds identified here do not inhibit bacterial growth and may therefore limit the development of antibiotic resistance if developed for use as novel treatments for chronic *P. aeruginosa* infections.

## Results and Discussion

### High throughput screening for repressors of EPS gene expression

A HTS for compounds that repress expression from a *pelB::lux* reporter was performed in 384-well microplate format using the Canadian Chemical Biology Network drug library containing 31,096 small molecule compounds. The *P. aeruginosa pelB::lux* reporter was grown in defined BM2 medium with limiting 20 μM magnesium (Mg^2+^), which we have identified previously as a growth condition that promotes biofilm formation, due to increased *pel/psl* expression and increased EPS production [4]. In the primary HTS screen, we tested the ability of compounds at 10 μM to reduce *pel* gene expression. Gene expression, in counts per second (CPS), was measured at a single time point (14 hours) in each well of 384-well microplates and normalized to the mean gene expression of each microplate. With this approach we were able to identify 163 compounds that reduced *pel* gene expression by at least 50% (Figure 1A). In a secondary screen of retesting the initial 163 hits, 14 compounds were identified that consistently repressed *pelB::lux* expression by 50% or greater, without any effect on growth. We next determined that the top 14 compounds also repressed a *pslA-lux* reporter by at least 50%. The structures to the 14 *pel* repressors are shown in Figure 2 and are described in Table 1.

**Figure 1.**
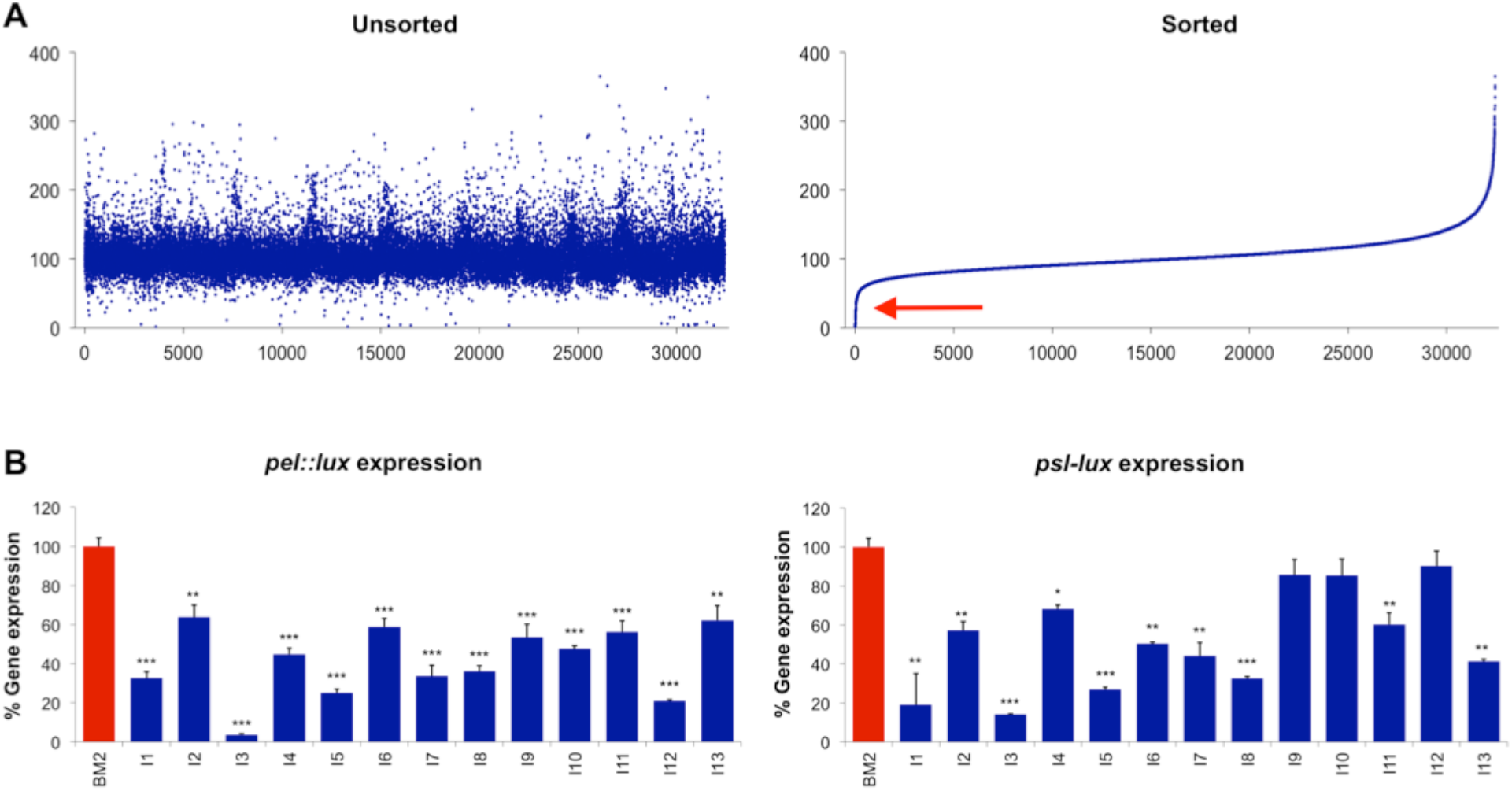
HTS identifies small molecules that reduce *pelB* and *psIA* expression. **(A)** Unsorted and sorted summary from a HTS showing the effect of each individual 31,096-small molecule on *pelB::lux* expression. Gene expression was measured at 14 hours and divided by the mean gene expression from each 384-well plate and represented as % gene expression. The primary screen identified 163 repressor “hits” (red arrow) that inhibited gene expression by 50% or more. **(B)** Reordered lead compounds identified in the HTS were retested for their ability to repress expression of *pelB::lux* and *pslA-lux* transcriptional reporters. Gene expression was measured throughout 18 hours and represented as area under the curve. Values shown are the average from triplicate values with standard deviation, n=4. Significant repression is indicated: *(p<0.05), **(p<0.01) and ***(p<0.001).

**Table 1:**
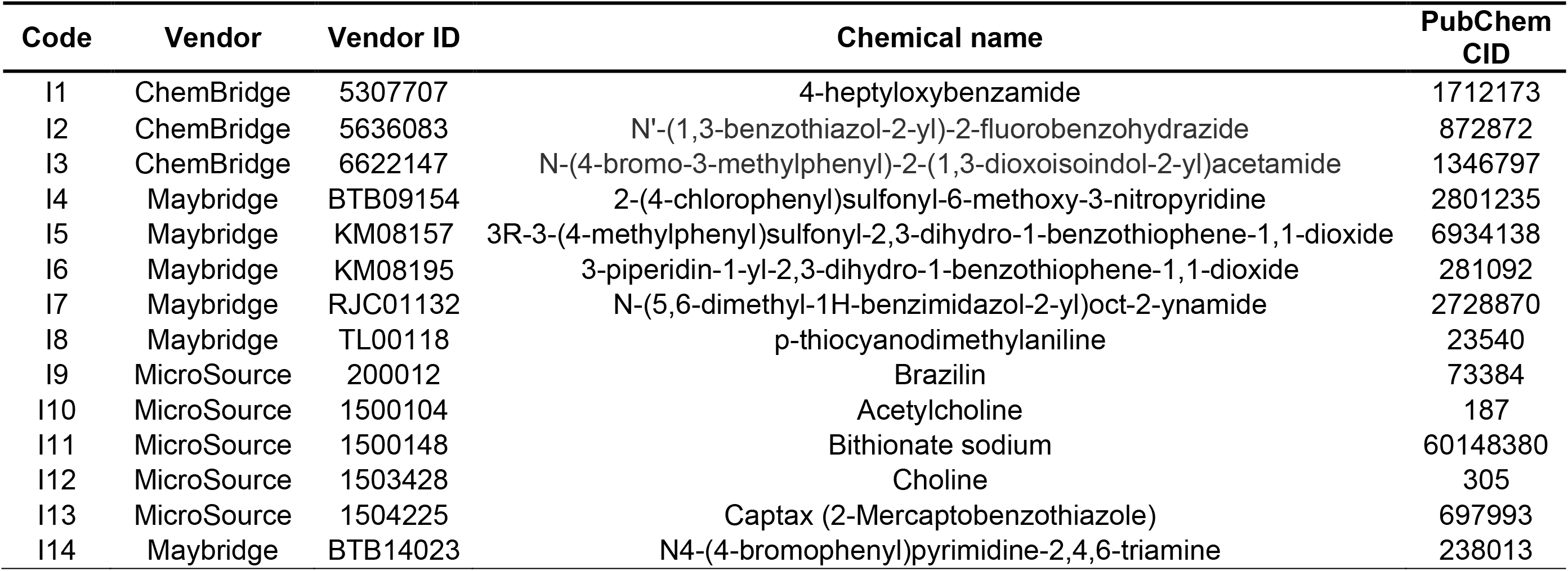
Molecular nomenclature of the *pel/psl* gene inhibitors identified in the HTS.

There were an additional 26 compounds that acted as *pelB::lux* repressors but that also had bactericidal or bacteriostatic properties (Table S2). These antimicrobial compounds could be separated into three different groups: small molecules with known antibiotic/disinfectant properties (group A), characterized non-antimicrobial molecules (group B) and uncharacterized molecules (group C) (Table S2). The identification of this panel of compounds highlights the ability of our approach to identify *pel* inhibitors, as well as antimicrobial molecules. It is noteworthy to mention that among these uncharacterized molecules (group C), two compounds (SPB07211 and KM07965) were recently identified in a whole-cell based HTS for small molecules that inhibit *Burkholderia cenocepacia* growth [25], and one small molecule (KM06346) was identified in a screen for nonspecific inhibitors of DNA repair enzymes (AddAB helicase nuclease) in *Escherichia coli* [26]. Nevertheless, all these growth inhibitor compounds were removed from the study.

To confirm the gene repression ability of these compounds, 13 of the 14 molecules (I1-I13) were reordered and retested. The 13 gene inhibitor compounds show a 30-90% repression of *pelB::lux* reporter gene expression (Figure 1B), in comparison to the same reporter strain cultured in biofilm inducing conditions alone. Additionally, most of the compounds were also able to reduce expression of *pslA-lux* reporter by 10-85% (Figure 1B), showing an effect over both EPS gene clusters. To determine if any of the compounds had a nonspecific effect on *lux* (luminescence), we grew a control *lux* reporter to the16S ribosomal RNA genes (PAO1::p16Slux) [27] in the presence of 10 μM compounds and measured gene expression throughout growth. There were minimal effects on 16S *lux* expression for most of the molecules, however one compound was a strong *lux* repressor (I3) and interestingly, one compound was a *lux* activator (I2) (Figure S1A). Therefore, we repeated the gene expression experiments with the *pelB::lux* and *pslA-lux* reporters and controlled for the compound effect on PAO1 *::p16Slux*, and again demonstrated that most compounds were repressors of both the *pel* (12/13) and *psl* (7/13) genes, relative to their effects on 16S expression (Figure S1B).

### Pel/Psl repressors reduce EPS secretion and biofilm formation under microplate and flow conditions

Considering the essential role of Pel and Psl for matrix formation and biofilm development, we initially assessed the ability of the identified gene repressors to reduce EPS synthesis in the wild type PAO1 strain. EPS secretion was quantitated by the standard congo red (CR) binding assay [28]. As predicted, most of the *pel/psl* repressors caused significant reduction in EPS secretion (Figure 3A). Given that the small molecules efficiently reduce EPS secretion, we next wanted to investigate the compounds ability to reduce biofilm formation.

**Figure 2.**
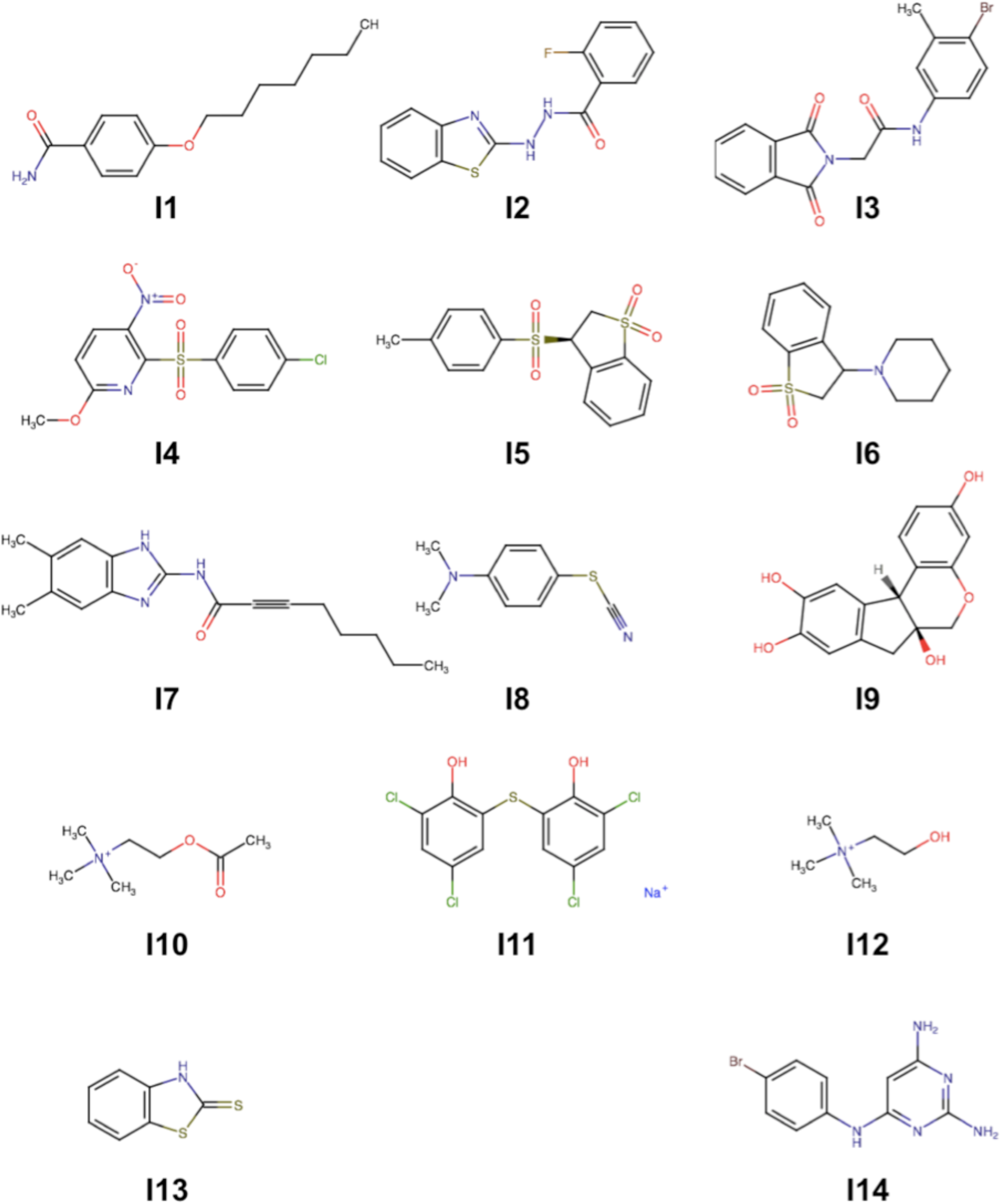
Molecular structures of the *pel/psl* gene expression inhibitors identified in the HTS. Inhibitor (I) compounds are labeled 1 through 14. For more details see Table 1.

**Figure 3.**
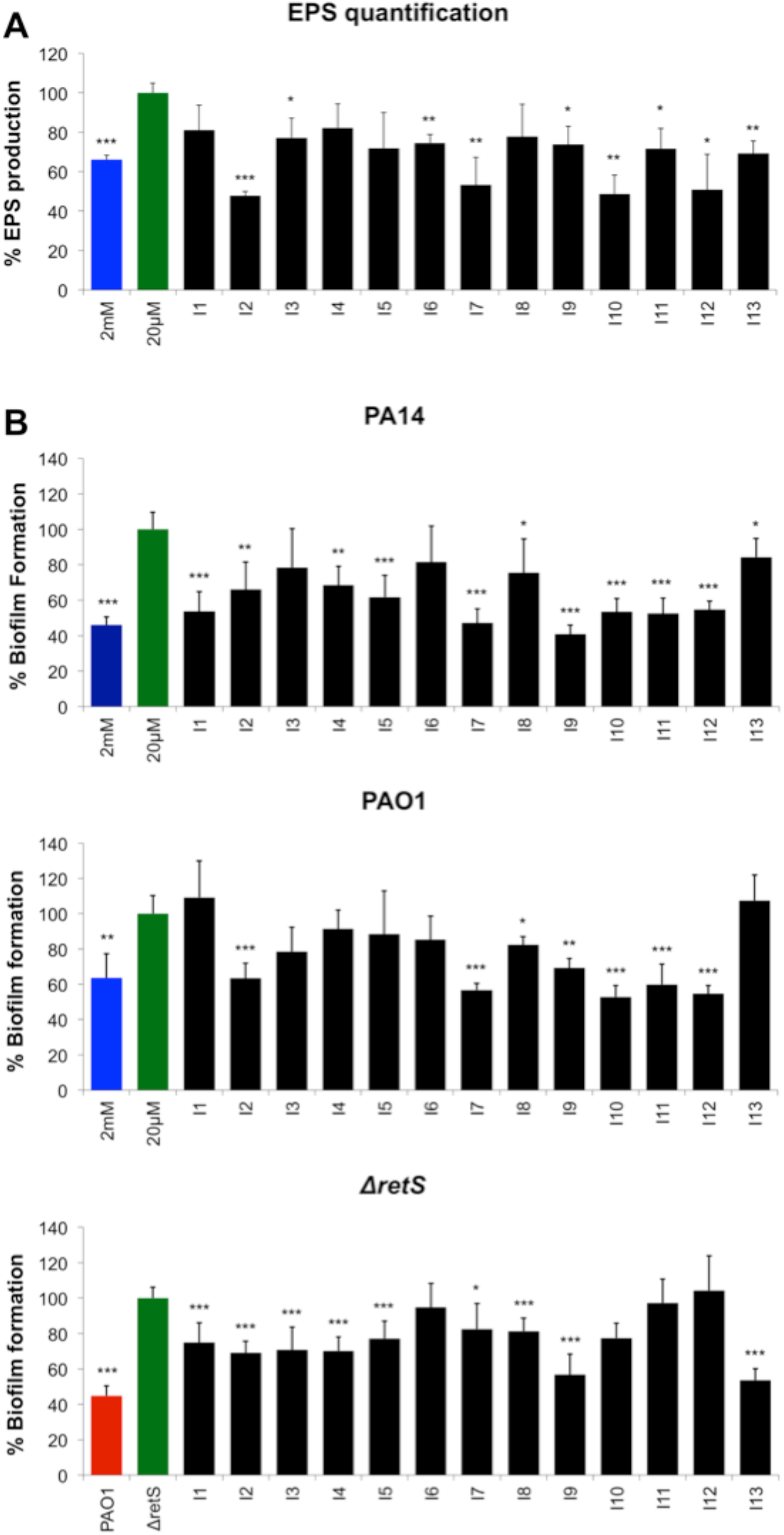
Pel/Psl repressor compounds reduce EPS production and biofilm formation *in vitro*. **(A)** Congo red (CR) was used to stain and quantitate secreted EPS after treatment with the 13 inhibitor compounds, n=3. **(B)** Crystal violent (CV) staining of biofilms formed in microplates after treatment of PA14, PAO1 and *retS::lux* biofilms. Black bars of compound-treated biofilms were compared to green bars of strains grown in biofilm-inducing condition alone. Blue (PA14 and PAO1) and red (PAO1) bars are negative controls of biofilms formed in repressing conditions. Values shown are the mean of 6 replicates plus standard deviation, n=3. Significant repression in biofilm formation is indicated: *(p<0.05), **(p<0.01) and ***(p<0.001).

*P. aeruginosa* biofilms were cultivated in 96-well microplates in the absence or presence of each of the top *pel/psl* repressor compounds and quantitated using crystal violet (CV) staining [29]. The *pel/psl* repressors caused significant reduction in the formation of microplate biofilms (Figure 3B). To better observe their antibiofilm effects in different strains of *P. aeruginosa*, we selected three different strains that differ in their ability to produce the two EPS molecules. The PA14 strain, due to a 3-gene deletion in the *psl* cluster, is only able to produce and secrete Pel, while PAO1 is able to produce both EPS molecules [10]. The majority of *pel/psl* repressors (11/13) promoted a significant reduction in biofilm formation in PA14 (Figure 3B), and most were effective (7/13) in reducing total biofilm biomass produced by PAO1 strain (Figure 3B). We also tested their antibiofilm activity against the *retS::lux* mutant, which is known to overproduce both the Pel/Psl and therefore have a hyperbioflm phenotype [30], similar to the small colony variants that arise in biofilms and in the CF lung [2]. Interestingly, 10/13 compounds were also effective in reducing biofilm formation in the insertional mutant *retS::lux* strain (Figure 3B). We were curious to assess whether there may be synergistic effects of combining the compounds; therefore, we tested the combination of 2 or 3 inhibitor compounds (at a total concentration of 10 μM) that we identified as strong repressors of *pelB::lux* expression (data not shown). We observed that some combinations were synergistic and resulted in greater degrees of biofilm inhibition than individual compounds (Figure S2A-C).

We next wanted to assess whether our compounds could inhibit the formation of biofilms in continuous flow systems, which tends to better mimic natural biofilms due to hydrodynamic influences [31, 32]. We cultivated and quantitated green fluorescent protein (*gfp*)-tagged PA14 and PAO1 biofilms in the BioFlux biofilm device (Figure 4A). We selected 9 individual antibiofilm compounds and 3 mixtures that showed strong biofilm inhibition in microplates. The majority of individual compounds and the combination treatments significantly reduced biofilm formation against PAO1-*gfp* and PA14-gfp, reducing both the depth (Figure 4B) and total coverage (Figure 4C) of biofilms grown in the BioFlux channel walls. Consistent with the CV biofilm assays (Figure S2), all compound mixtures tested had greater effects on reducing biofilms than individual compounds alone (Figure 4).

**Figure 4:**
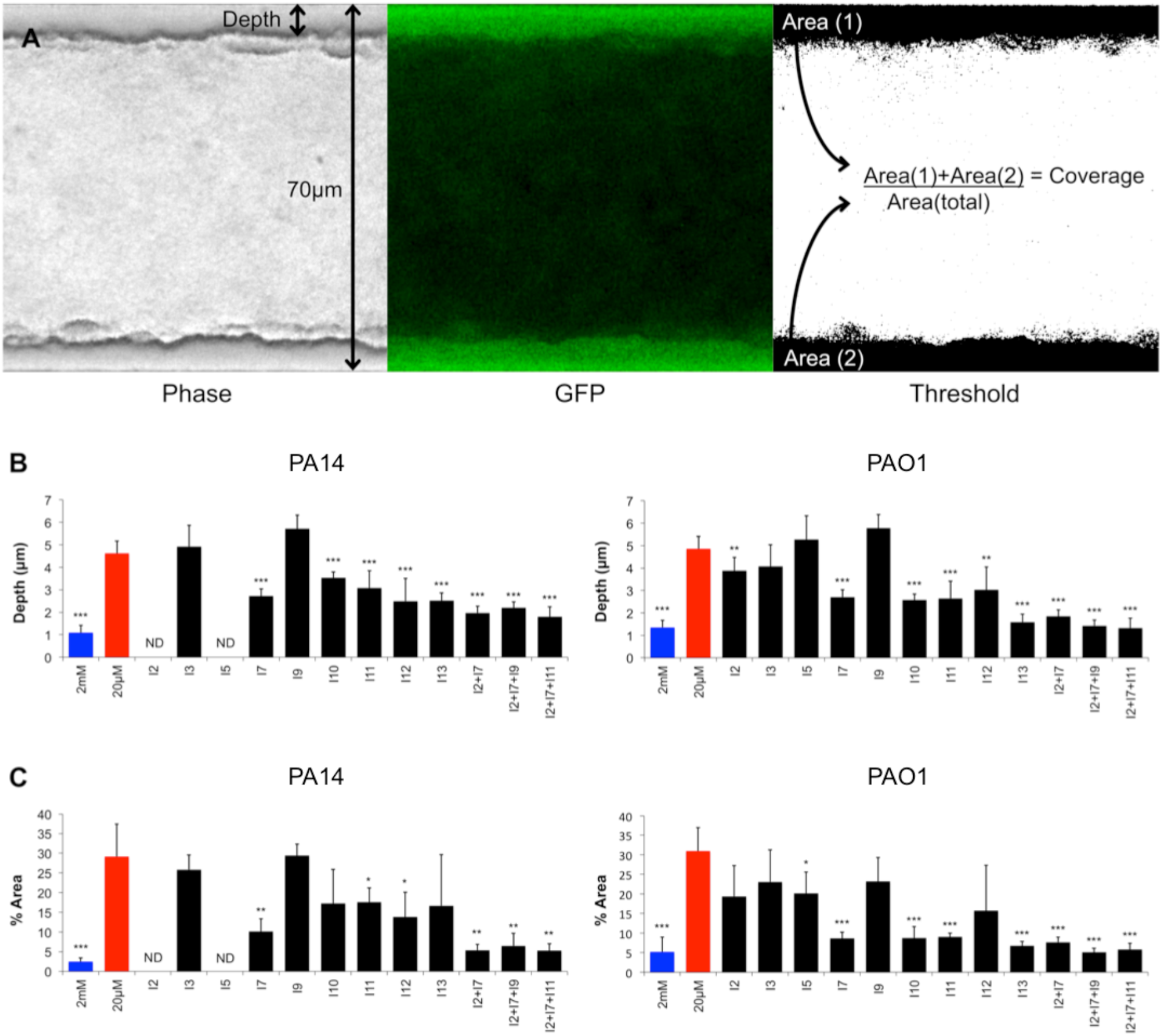
EPS inhibitors reduce biofilms formed under flow conditions. **(A)** The images shown of biofilms formed along the walls of the chambers in the BioFlux device are from phage contrast (left), GFP (middle) and the GFP image that was threshold adjusted to isolate the biofilms adhered to the channel walls (right). The phase contrast images were used for calculating the **(B)** biofilm depth, and the GFP images were used for calculating **(C)** the total biofilm coverage in the chamber. Black bars of compound-treated biofilms were compared to strains grown in biofilm-inducing (green) or repressing (blue) conditions. All values shown are the mean of triplicate samples plus standard deviation, n=2. Significant repression in biofilm coverage or depth is indicated: *(p<0.05), **(p<0.01) and ***(p<0.001).

### Pel/Psl repressors reduce mucoid biofilm formation and non-mucoid biofilms under anaerobic conditions

It is known that the chronically infected, mucus-filled airways of the CF lung lead to the selection of mucoid and alginate-overproducing *P. aeruginosa* [19, 33, 34]. Mucoid strains promote long-term survival in the CF lung, by promoting aggregation, antibiotic resistance and increased resistance to host immune response such as oxidative stress [19, 34]. Given the abundance of mucoid isolates in the CF lung, and the necessity of both intact *pel* and *psl* gene operons for biofilm formation and fitness of mucoid strains [35], we wanted to determine if our antibiofilm compounds could also reduce biofilm formation in a mucoid variant. Biofilms of strain PDO300, a *ΔmucA22* variant of PAO1 [36], were cultivated in *pel/psl* non-inducing conditions. BM2 containing 2mM Mg^2+^ was used to minimize the contribution of Psl/Pel production in this experiment. Six antibiofilm compounds present at 10 μM promoted a modest but significant reduction in biofilm formation for the mucoid strain (Figure 5A). Interestingly, two of the Psl/Pel antibiofilm compounds had a biofilm promoting effect on the mucoid variant of PAO1 (Figure 5A).

**Figure 5.**
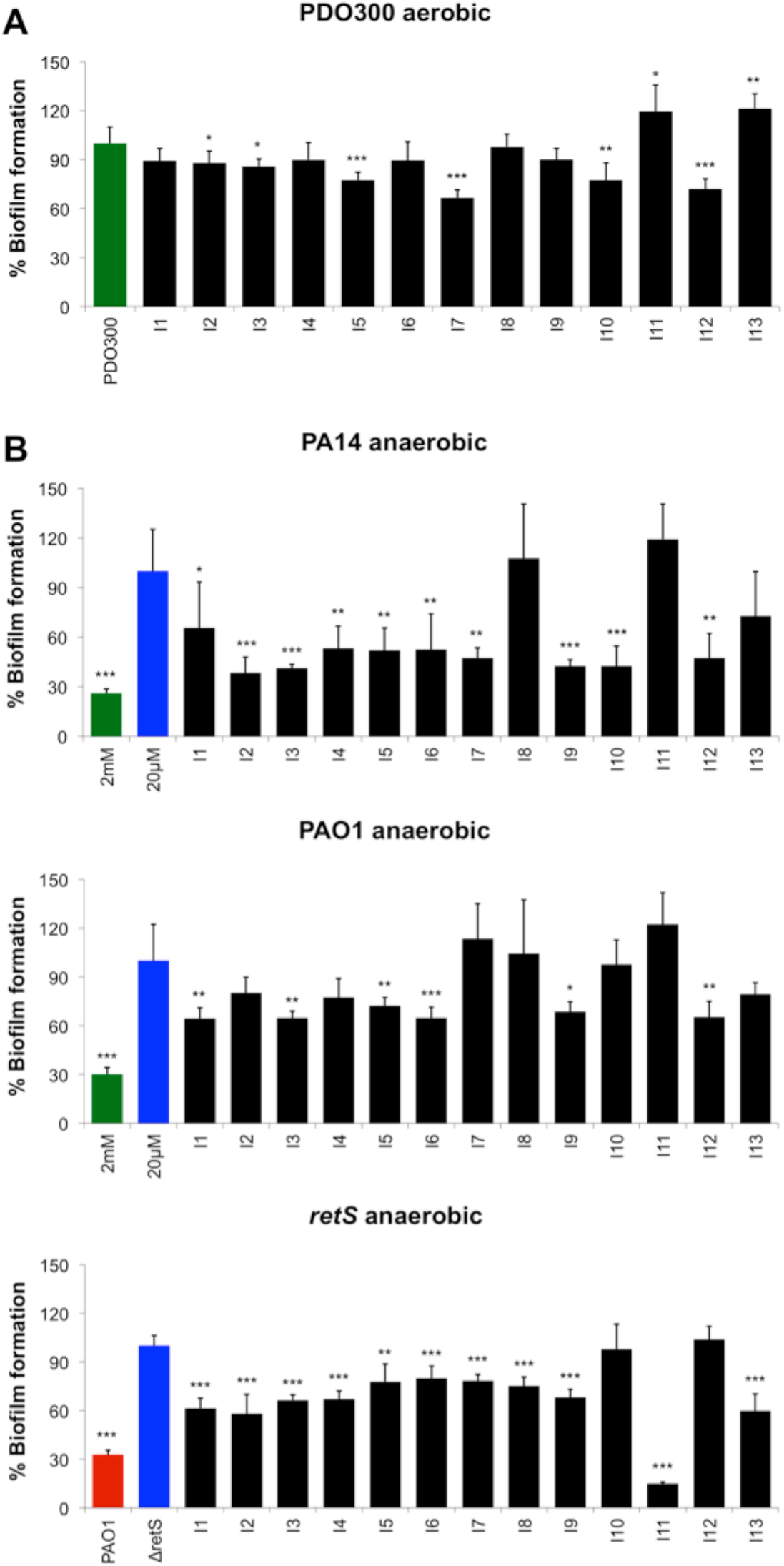
Small molecule EPS inhibitors reduce biofilms formed by mucoid strains and non-mucoid strains under anaerobic conditions. **(A)** Crystal violent (CV) staining of mucoid PDO300 (*ΔmucA22*) biofilms in aerobic conditions. Black bars of compound treated biofilms were compared to green bar of mucoid biofilms grown in EPS-repressing condition. **(B)** Crystal violent (CV) staining of anaerobic biofilms formed in microplates after treatment of PA14, PAO1 and *retS::lux* strains. Black bars of compound-treated biofilms were compared to green bars of strains grown in biofilm-inducing condition alone. Blue (PA14 and PAO1) and red (PAO1) bars for negative controls of biofilms formed in repressing conditions. ND, not determined. All values shown are the mean of 6 replicates plus standard deviation, n=3. Significant repression in biofilm formation is indicated: *(p<0.05), **(p<0.01) and ***(p<0.001).

Previous reports have shown that the CF mucus plug environment consists of both aerobic and anaerobic microenvironments [37]. Therefore, we tested our compounds for their ability to reduce biofilm formation under anaerobic conditions. The antibiofilm compound treatments (10/13) were able to promote a significant reduction in biofilm formation in the PA14 strain under anaerobic conditions, while 6/13 compounds were effective against the PAO1 strain (Figure 5B). Importantly, 10/13 compounds also reduced biofilm formation in the hyperbiofilm forming mutant *retS::lux* strain under anaerobic conditions (Figure 5B). Although most of the compounds showed a similar trend in biofilm repression for both aerobic and anaerobic conditions, compound I6 showed antibiofilm properties only under anaerobic growth. Additionally, we were unable to determine biofilm inhibition by compound I11, as it was inhibiting bacteria growth under anaerobic conditions for the three strains tested (data not shown).

### Pel and Psl are required for PAO1 full virulence in the *C. elegans* infection model

Although Pel and Psl are well appreciated for their role in biofilm formation, their contribution to virulence is less understood. *P. aeruginosa* is known to colonize the intestinal lumen of *C. elegans* and cause severe alterations in its morphology [38]. Furthermore, *P. aeruginosa* forms clumps in the nematode gut, surrounded by an uncharacterized extracellular matrix [38]. Therefore, to determine whether Pel and/or Psl contribute to bacterial virulence we utilized the nematode infection model. We assessed the nematode feeding preference and slow killing assays when *C. elegans* were fed *P. aeruginosa* possessing mutations in the *pel* and *psl* gene clusters. Initially we used the feeding preference assay, where the nematodes are given an option of test strains within a grid of 48 colonies [39]. As feeding is observed until colony disappearance, strains that are eaten preferentially were also shown to be less virulent in the slow killing assays [39]. While single knockouts in either the *pel* or *psl* genes did not alter the nematode feeding behavior, a double *Δpel/psl* mutant was preferentially eaten by *C. elegans*, when given the choice of between *Δpel/psl* and PAO1 (Figure 6A). As a control experiment, the laboratory food source *E. coli* OP50 also served as a preferential food source to PAO1 (Figure 6A). None of the mutant strains tested in the feeding preference assay showed a growth defects in SK media (data not shown). This result suggests a virulence role for Pel and Psl *in vivo*.

**Figure 6.**
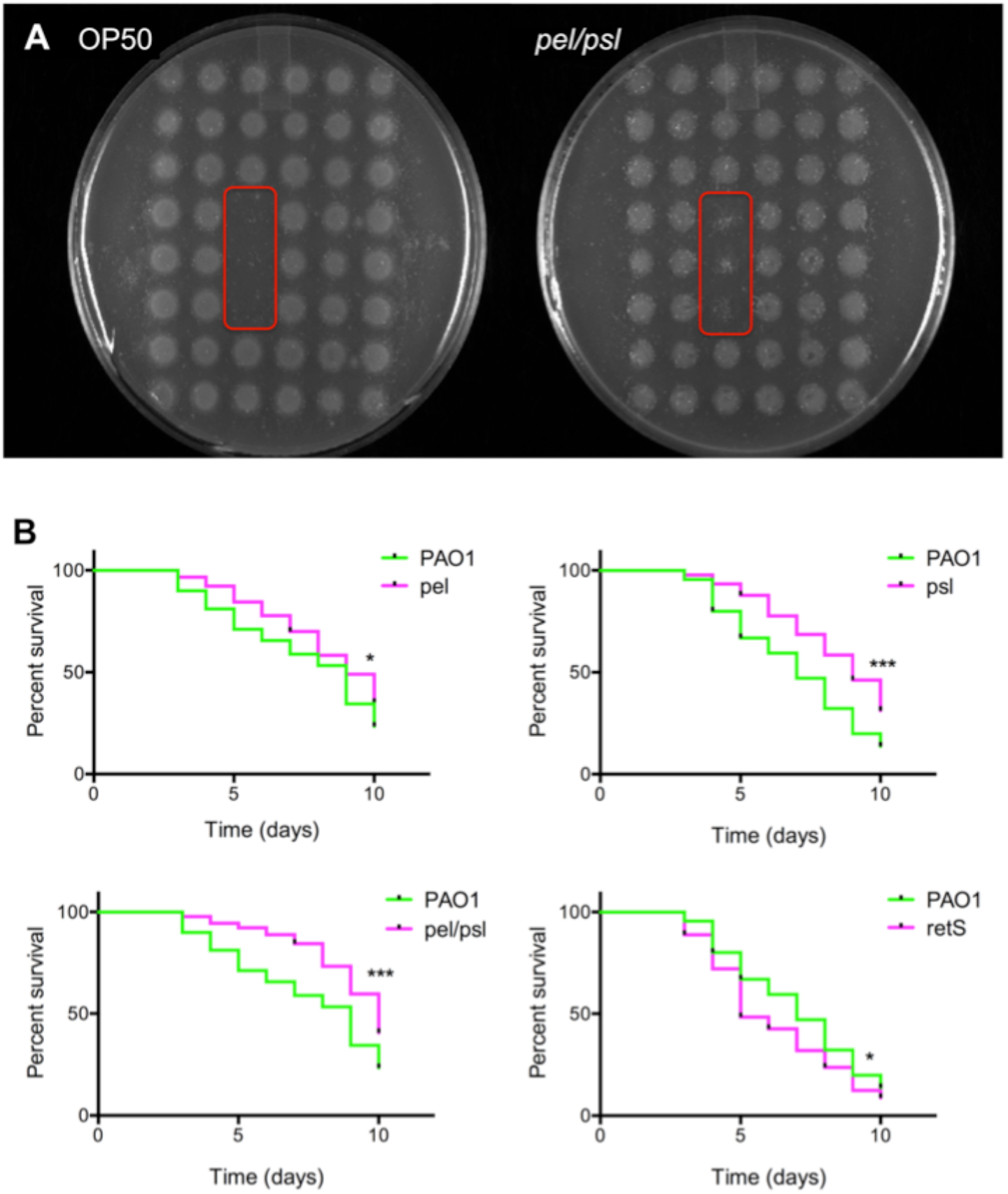
The Pel and Psl exopolysaccharides are required for full virulence in the *C. elegans* infection models. **(A)** The feeding preference assay indicates that the *pel/psl* double mutant is a preferred food source and was eaten to completion before PAO1. Test strains were embedded in triplicate spots (red box) within a grid of 6x8 positions of wild type PAO1, n=3. The *E. coli* OP50 strain was used as a positive control of preferred food source known to have reduced virulence (see figure S3). **(B)** Slow killing curves of nematodes fed individual strains of either PAO1 wild type, *pel, psl* or *pel/psl* mutants. The hyperbiofilm forming *retS::lux* mutant was also tested for virulence relative to the PAO1. Kaplan-Meyer curves with % survival represent three independent experiments (n=30) where the total number of worms equals 90. Significant differences in nematode survival are indicated: *(p<0.05), **(p<0.01) and ***(p<0.001).

To further investigate the role of EPS in virulence, we conducted slow killing assays [39, 40], in which nematodes are given a single bacterial food source, and worms survival is monitored over 10 days. In addition to the single and double *pel/psl* mutant panel, we also tested the Pel/Psl hyperproducing *retS::lux* mutant. In the slow killing assay, both the single *pel* and *psl* mutants, as well as the double *Δpel/psl* were less virulent and resulted in increased nematode survival throughout 10 days (Figure 6B). Interestingly, the *retS::lux* mutant, known to overproduce both Pel and Psl, demonstrated an increased virulence in the slow killing assay (Figure 6B). Although this effect in the *retS::lux* mutant may be due to other pleiotropic effects of mutation in this regulatory protein [30], taken together, these observations indicate that both the Pel and Psl are required for full virulence of *P. aeruginosa* in killing *C. elegans*.

### Antibiofilm molecules also have antivirulence activity

Since the Pel/Psl are required for *C. elegans* full virulence (Figure 6), we hypothesized that the antibiofilm compounds identified in the HTS would also reduce virulence of the wild type PAO1. For the slow killing assay, PAO1 was inoculated as a lawn on agar plates that also included 10 μM of the antibiofilm compounds. Next, L4 stage nematodes were transferred to the plate containing compound-treated PAO1 food sources and survival was monitored over time. Although no significant killing effects were observed for the majority of the compounds tested (Figure S4), compounds I7, I9, I10 and I11 caused a significant reduction in PAO1 virulence after 10 days of feeding on compound-treated bacteria (Figure 7). This increase in nematode survival was comparable to the effect of inactivating the Pel and/or Psl EPS (Figure 6B), suggesting that these small molecules demonstrate antivirulence activity for *P. aeruginosa* due to repression of EPS synthesis.

**Figure 7.**
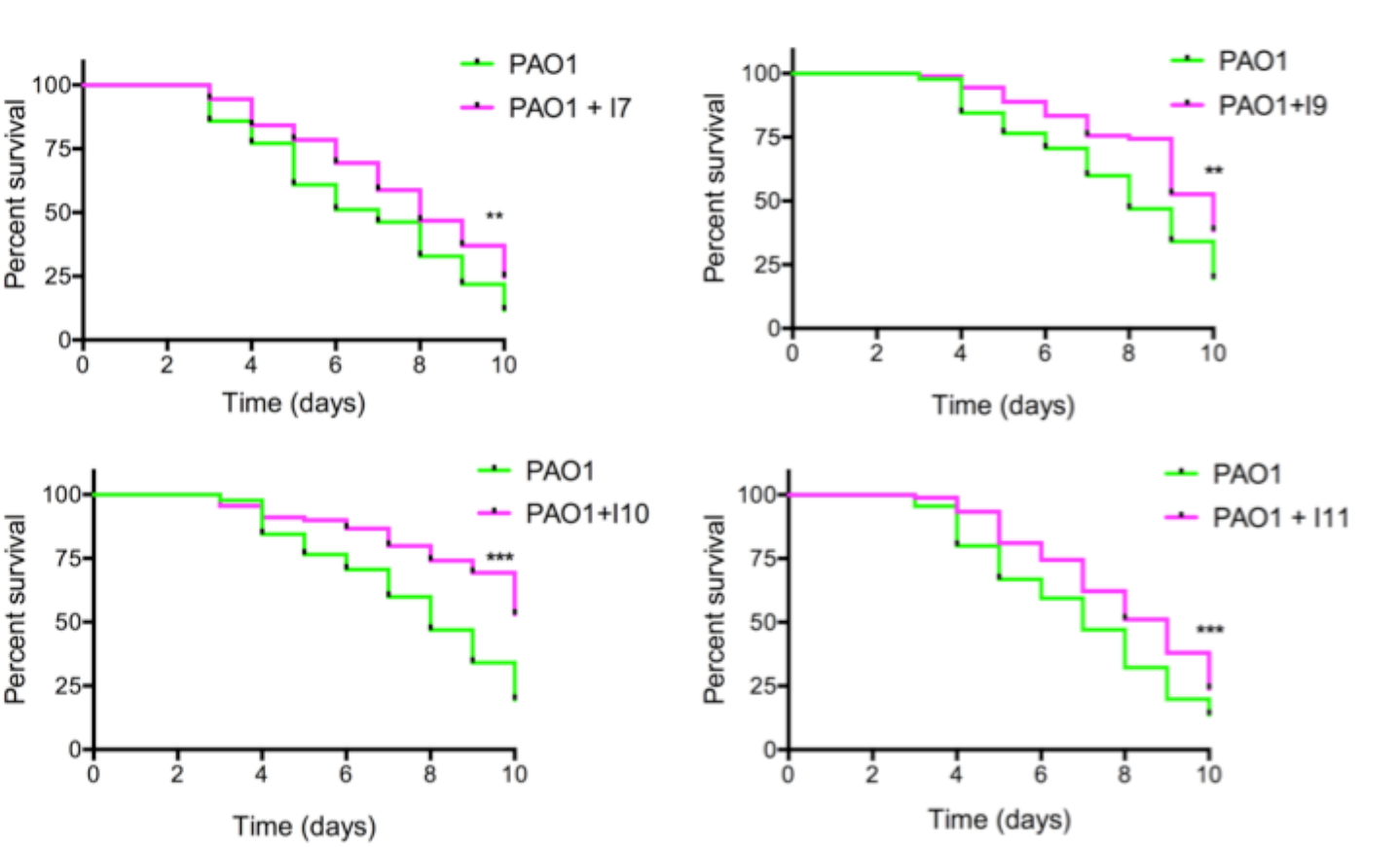
Antibiofilm compounds have antivirulence activity in the *C. elegans* slow killing infection model. Nematodes were fed with antibiofilm compound-treated PAO1 and monitored for increased survival. Kaplan-Meyer curves with % survival represent three independent experiments (n=30) where the total number of worms equals 90. Significant differences in nematode survival are indicated: **(p<0.01) and ***(p<0.001).

### Antibiofilm small molecules are synergistic with antibiotic killing against PAO1 biofilms

Since biofilms are more antibiotic tolerant than planktonic cells, new treatments are needed that can both reduce biofilm and increase antibiotic susceptibility [22]. Peg-adhered PAO1 biofilms were cultivated in the absence or presence of our lead antibiofilm compounds for 24 hours, and then challenged with a panel of 5 different antimicrobials that target *P. aeruginosa* growth by different mechanisms of action. Polymyxins (colistin and polymyxin B) disrupt membrane integrity, aminoglycosides (tobramycin and gentamicin) inhibit protein synthesis and fluoroquinolones (ciprofloxacin) block DNA replication. It is noteworthy to acknowledge that Col and Tm are two clinically important antimicrobial therapies used to treat *P. aeruginosa* infections in CF patients [41, 42].

Biofilms were first cultivated in a gradient of increasing concentrations of our antibiofilm compounds and then challenged with sub-biofilm eradication concentrations of the antimicrobials. We selected the four molecules that demonstrated both antibiofilm and antivirulence activities. Compounds I7, I10 and I11 increased the PAO1 biofilm susceptibility to all tested antimicrobials by reducing the viable cell counts between 10 and 10,000 fold (Figure 8). Compound I9 only increased biofilm susceptibility to Ci and PB and compound I11 often caused a dose-dependent effect on increasing biofilm killing as the concentration of antibiofilm compound increased from 5 to 20 μM (Figure 8). Most compounds had the strongest potency in increasing biofilm sensitivity at 5 and 10 μM concentrations (Figure 8). In addition to their antibiofilm and antivirulence properties, these small molecules are also synergistic with antibiotics due to their ability to reduce overall biomass, which likely improves antibiotic access and their ability to kill bacteria.

**Figure 8.**
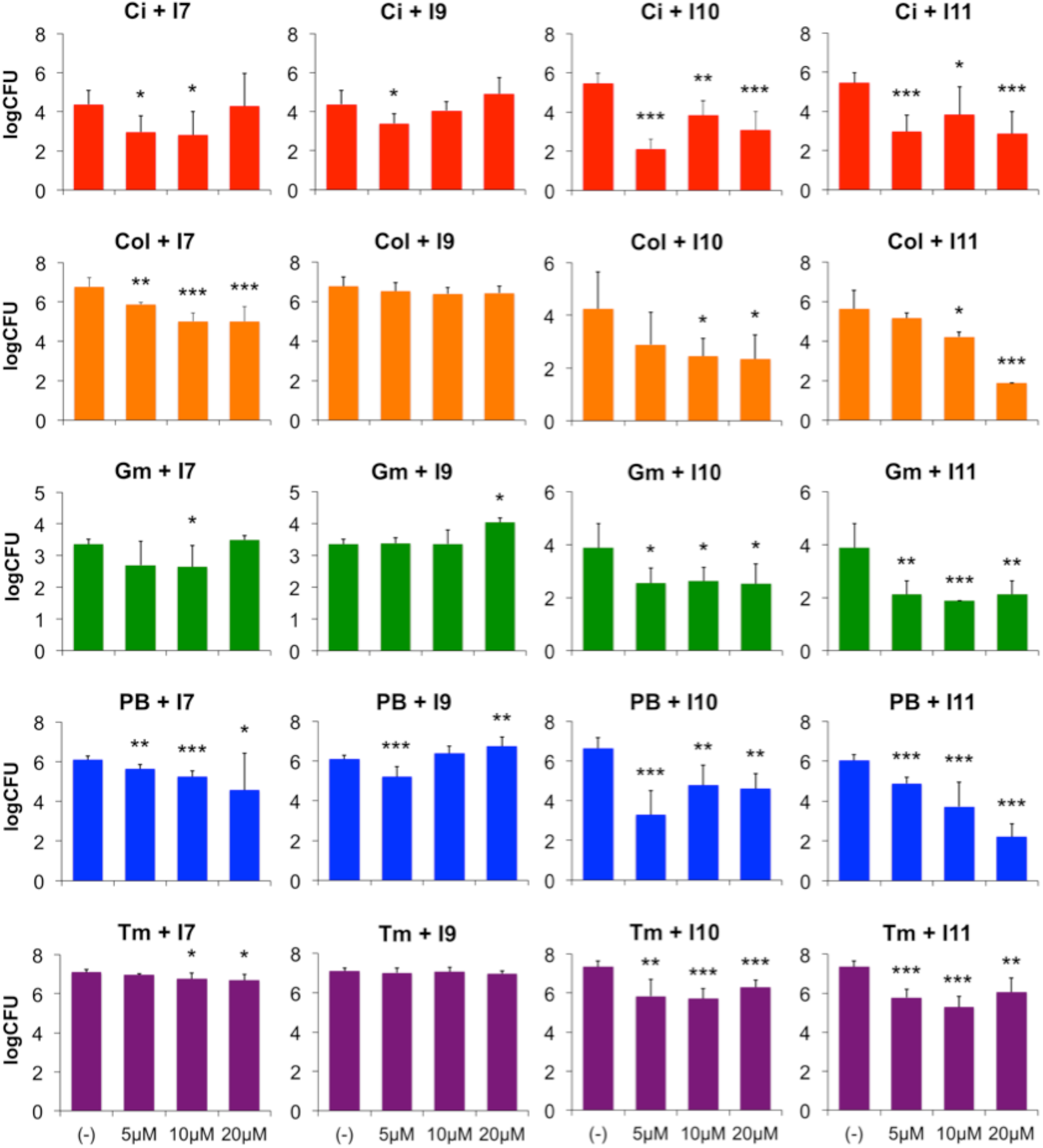
Antibiofilm compounds have synergistic effects when used in combination with conventional antibiotics. Biofilms were formed in the absence or presence of antibiofilm compounds I7, I9, I10 and I11, and then treated with sub-eradication concentrations of ciprofloxacin (Ci, 2.5 μg/ml), colistin (Col, 25 μg/ml), gentamicin (Gm, 6.5 μg/ml), polymyxin B (PB, 25 μg/ml) or tobramycin (Tm, 1 μg/ml) antibiotics. CFU/ml counts of surviving bacteria were determined from peg-adhered biofilms after antibiotic treatment. Antibiofilm compounds were added in increasing concentrations ranging from 0-20 μM. Values shown are the mean of triplicate samples plus standard deviation, n=2.

### The Gac/Rsm regulatory network as a possible target

We speculated that the lead antibiofilm compounds might be acting on one of the regulatory components of the intricate signaling network that controls the expression and production of Pel and Psl (Figure 9A). Most attention is paid to the role of the GacAS two-component system and the *rsmY* and *rsmZ* regulatory RNAs in regulating the mRNA stability by RsmA and production of Pel and Psl [43–45]. In addition, the GacAS pathway can be repressed or activated by additional orphan sensors, RetS and LadS, respectively [30, 46]. Other transcriptional regulators include the *psl* activator RpoS [45] and the repressor/activator FleQ [47] (Figure 9A). RetS was originally described as a transcriptional repressor of *pel* and *psl* using microarray analysis [30]. The mechanism of RetS transcriptional control of *pel/psl* is not understood, given the lack of a known cognate response regulator. We previously reported that the PhoPQ two component system directly represses the *retS* gene, which may account for the robust biofilm phenotype under the PhoPQ-inducing conditions of growth in limiting Mg^2+^ [4]. Repressing the biofilm inhibitor RetS, may lead to increased biofilm through either transcriptional or post-transcriptional control of Pel/Psl.

**Figure 9.**
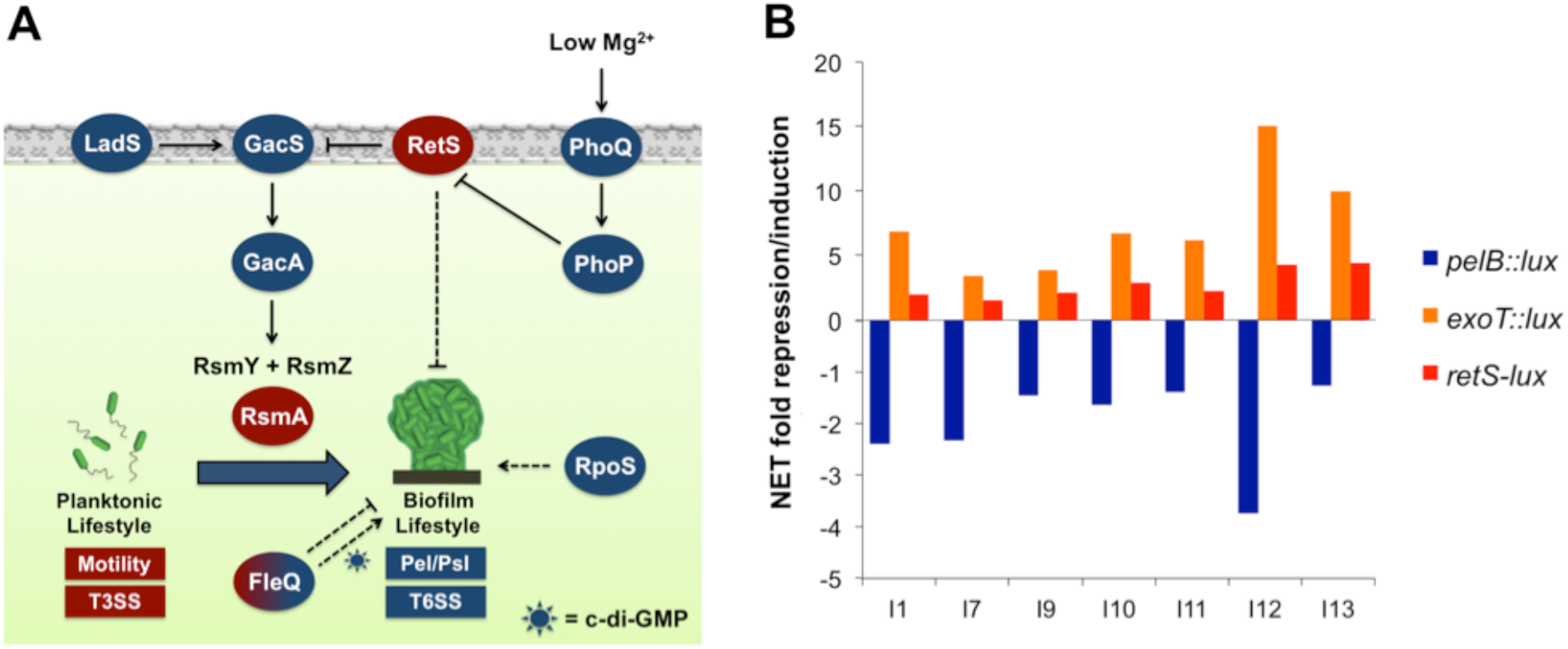
The central regulatory systems controlling Pel/Psl EPS production and biofilm formation. **(A)** The Gac/Rsm regulatory pathway in *P. aeruginosa* is the central pathway that post-transcriptionally controls the switch to a biofilm lifestyle. Whether activated by LadS, or repressed by RetS, both additional inner membrane orphan sensors, the Gac/Rsm pathway promotes stability of the Pel and Psl transcripts and ultimately EPS production and biofilm formation (solid lines). This pathway inversely controls the T3SS, largely thought to be an acute virulence factor and not required for chronic infections caused by biofilms. RetS also contributes to the transcriptional repression of *pel* and *psl*, along with the transcriptional activators RpoS and FleQ (dashed lines). FleQ can function as a repressor or activator of the *pel* operon, depending on the presence of cyclic-di-GMP. The Mg^2+^ sensing PhoPQ two-component system directly represses *retS*, which in turn induces biofilm formation. Positive regulators of the biofilm lifestyle are represented in blue, negative regulators in red. **(B)** Gene expression profiles of *pelB, retS and exoT* after treatment with select antibiofilm compounds. Bars represent the average net fold induction (positive) or repression (negative) from triplicate samples of the target genes relative to the untreated condition.

A central feature of the Gac/Rsm pathway is that when biofilm production is promoted, the type III secretion system (T3SS) is repressed [30]. To determine if these antibiofilm compounds potentially act on the Gac/Rsm pathway, we monitored the expression profiles of genes normally repressed under biofilm promoting conditions of limiting Mg^2+^, which includes the *retS* sensor and the *exoT* T3SS effector [4, 30].

Several of the antibiofilm compounds caused the induction of *retS* biofilm repressor and *exoT*, along with the simultaneous repression of *pel* (Figure 9B), which is the opposite pattern of expression in the biofilm mode of growth. By reversing the expression of genes controlled by the Gac/Rsm pathway, the compounds identified in this study may act somewhere in this regulatory pathway. Some compounds were ineffective in reducing biofilms formed by the *retS* mutant (I10, I11), suggesting that RetS may be a possible drug target in these cases. However, given the large number of potential protein targets (Figure 9A), further work is needed to confirm their possible mechanism of action.

## Conclusion

Antivirulence compounds have been described for *P. aeruginosa* that target the quorum sensing Las, Rhl and Pqs systems [48]. Here we describe the *P. aeruginosa* EPS biosynthesis genes as a new target for the identification of antivirulence compounds. Biofilm formation is an important focus for new antimicrobials given the universal and conserved process of forming a biofilm, and the diverse protective advantages of cells enmeshed in an extracellular matrix. We identified compounds that repress the expression of the *pel* and *psl* EPS genes in *P. aeruginosa* and hypothesized that this effect would lead to biofilm defective phenotypes. Next we illustrated that EPS production is required for *P. aeruginosa* virulence in nematode infection (Figure 6), and 4 of the 13 *pel/psl* inhibitors reduced the virulence of PAO1 in the slow killing infection model for *C. elegans* (Figure 7). Further, by reducing EPS synthesis and biofilm formation (Figure 3), the antibiofilm molecules also demonstrated synergistic activity when combined with antibiotic treatment (Figure 8). The antibiotic synergies were seen across multiple antibiotic classes, including antibiotics previously shown to be affected by the production of Pel/Psl EPS [8, 15, 17].

Future work will focus on identifying the mechanism of action of the molecules found in this study. Some of the compounds appear to act on the Gac/Rsm pathway, as they reverse the pattern of target gene expression. We used the biofilm promoting conditions of growth in limiting Mg^2+^ concentration, which increases EPS production [4] and represses expression of the T3SS [49]. Several of the antibiofilm compounds in this study reversed that pattern, by repressing EPS biosynthesis and inducing the T3SS, as well as induce the RetS biofilm repressor (Figure 9B). Here we identified diverse structural compounds that possess antibiofilm activity, some of which share common structural features. Compounds I5 and I6 share a benzothiophene backbone, compounds I2 and I13 have a benzothiazole component, and I10 and I12 are acetylcholine and choline, respectively (Table S2). In support of our findings, choline analogs have been previously identified as antibiofilm agents against *P. aeruginosa* [50]. In summary, we have identified a panel of small molecules that represent a novel class of antivirulence antimicrobials, which could be developed for the treatment of chronic *P. aeruginosa* biofilm infections.

## Acknowledgments

The authors thank George Chaconas for providing the drug library, Elaine Goth-Birkigt for technical assistance with the BioFlux device and Joe Harrison for providing the plasmids for *PA14-gfp* construction. We also thank Joe McPhee, Tao Dong and Mike Wilton for helpful comments on the manuscript. Funding was provided by an NSERC Discovery Grant. EVTB was supported by the Beverly Phillips Rising Star and Cystic Fibrosis Canada Studentships and SL held the Westaim-ASRA Chair in Biofilm Research.

## Materials and Methods

### Bacterial strains and growth media

The strains and plasmids used in this study are listed in Table S1. Biofilms and planktonic cultures were grown in Basal Minimal Medium (BM2) at 37°C, containing excess or limiting Mg^2+^ concentrations. BM2 media prepared with 100 mM Hepes pH 7.0, 7mM (NH_4_)_2_SO_4_, 1.03 mM K_2_HPO_4_, 0.57 mM KH_2_PO4, 20 μM or 2 mM MgSO_4_, 10 μM FeSO_4_ and ion solution, containing 1.6 mM MnSO_4_.H_2_0, 14 mM ZnCl_2_, 4.7 mM H_3_BO_3_ and 0.7 mM CoCl_2_.6H_2_0. The media was supplemented with 20 mM sodium succinate as a carbon source for all assays. KNO_3_ (1%) was added to support bacterial growth under anaerobic conditions. GFP-tagged PA14 was prepared as previously described [51]. Briefly, the pBT270 GFP-encoding plasmid containing a site-specific integration mini-Tn7 vector was transformed in PA14, with the help of a pTNS2 plasmid. GFP-expressing colonies were selected for with Gm resistance (50 μg/ml). Stock solutions of ampicillin (50 mg/ml, AMRESCO), ciprofloxacin (2 mg/ml, BioChemika), colistin (10 mg/ml, Sigma), gentamicin (30 mg/ml, Sigma), polymyxin B (30 mg/ml, Sigma), and tobramycin (25 mg/ml, Sigma) were made in ultrapure water and stored at −20°C, and used as indicated.

### HTS and gene expression assays

For the HTS of the 31,096 small molecules in the Canadian Chemical Biology Network library, compounds were transferred from the stock 96-well plates (1 mM in dimethyl sulfoxide) to 384-well assay microplates with the help of plastic 96-pin transfer devices. All compounds were tested at a final concentration of ~10 μM. Plates were covered with an air-permeable membrane and incubated at 37°C for 14 hours, and gene expression in counts per second (CPS) was determined in a Wallac Victor^3^ luminescent plate reader (Perkin-Elmer). For the secondary screen, *pelB::lux* and *pslA-lux* reporters were grown in 384-well microplates in the presence of ~10 μM of the hit-compounds at 37°C in a Wallac Victor^3^ plate reader, CPS and optical density (growth, OD_600_) reads were taken every 60 minutes throughout growth. For all other gene expression assays, gene reporters were grown in 96-well microplates, in the presence of 10 μM of the reordered inhibitor compounds, and incubated at 37°C in a Wallac Victor^3^, with CPS and OD_600_ measurements taken every 20 minutes.

### EPS quantification

EPS was measured in the quantitative congo red (CR) binding assay, as previously described [28]. Cultures were grown in BM2 20 μM Mg^2+^ containing 10 μM of reordered compounds in 5 ml glass tubes for 24 hours at 37°C with shaking (150 rpm). For EPS quantification, CR was added at a final concentration of 40 μg/ml to 2 ml cultures and bound-dye was indirectly calculated by determination of remaining unbound dye still in solution [28].

### Biofilm cultivation and quantification

Unless otherwise indicated, all biofilms were cultivated in BM2 20 μM Mg^2+^ containing 10 μM of compounds in 96-well polystyrene microplates for 18 hours at 37°C with shaking (100 rpm). Anaerobic biofilms were cultivated inside an anaerobic chamber (GasPack System) for 48 hours. Biofilm inhibition was determined by crystal violet (CV) staining as previously described [29]. Continuous flow biofilms were cultivated in BioFlux device, in 48-well microplates, incubated for 18 hours at 37°C [31]. The biofilms formed under flow system were imaged with a Nikon Eclipse Ti inverted epifluorescence microscope in green fluorescence and phase contrast. Biofilm depth and coverage were determined using ImageJ processing and analysis in Java.

### *C. elegans* nematode infection models

The nematode feeding preference and slow killing assays were performed as previously described [39]. Briefly, for the feeding preference assay, 20 L4 stage hermaphrodite nematodes were transferred to a slow killing (SK) assay plate containing a pre-grown grid of 45 wild type colonies and 3 internal spots of the mutant strains to be tested (6x8 colonies). The plates were incubated at 25°C and observed twice a day until the disappearance of the initial bacterial colonies. For the slow killing assay, 30 L4 stage nematodes were transferred to a SK plate containing individual pre-grown PAO1 lawns of bacteria to be tested. Nematode survival was determined throughout 10 days by direct observation under a dissecting microscope. 25 μg/ml of 5-fluoro-2’-deoxyuridine (FUdR) was added to SK plates on the slow killing assay for the prevention of offspring development. All compounds were added at 10 μM in the SK agar before bacterial growth. For both *in vivo* assays, *E. coli* strain OP50 was used as a positive control to demonstrate preferred feeding behavior and reduced virulence for the nematodes. Nematodes were cultivated on nematode growth medium (NGM) plates containing a lawn of OP50 as food source.

### Antimicrobial synergy testing of biofilms

Biofilm susceptibility testing was performed as previously described [52], with minor modifications. Biofilms were grown for 24 hours on polystyrene pegs (NUNC-TSP) submerged in BM2 20 μM Mg^2+^ containing 5, 10 and 20 μM of select inhibitor compounds. Cultivated biofilms were rinsed in 0.9% saline (NaCl) and transferred to a plate containing sub-eradication concentrations of antimicrobials in 10% BM2 diluted in 0.9% saline and challenged for 24 hours. After challenge, biofilms were rinsed twice in 0.9% saline, and attached cells were DNase I treated (25 μg/ml, Sigma) for 30 minutes and sonicated for 10 minutes to detach. Cell viability was determined by serial dilution and direct counting as previously described [53].

### Statistical analysis

Statistical significance between populations was determined by paired two-tailed Student’s t-test. Log-rank test (Graph Pad Prism) was used for determination of significant differences in *C. elegans* survival. Data considered significant at level of p<0.05. Significance represented by * (p<0.05), ** (p<0.01) and *** (p<0.001).

---

**Figure S1:**
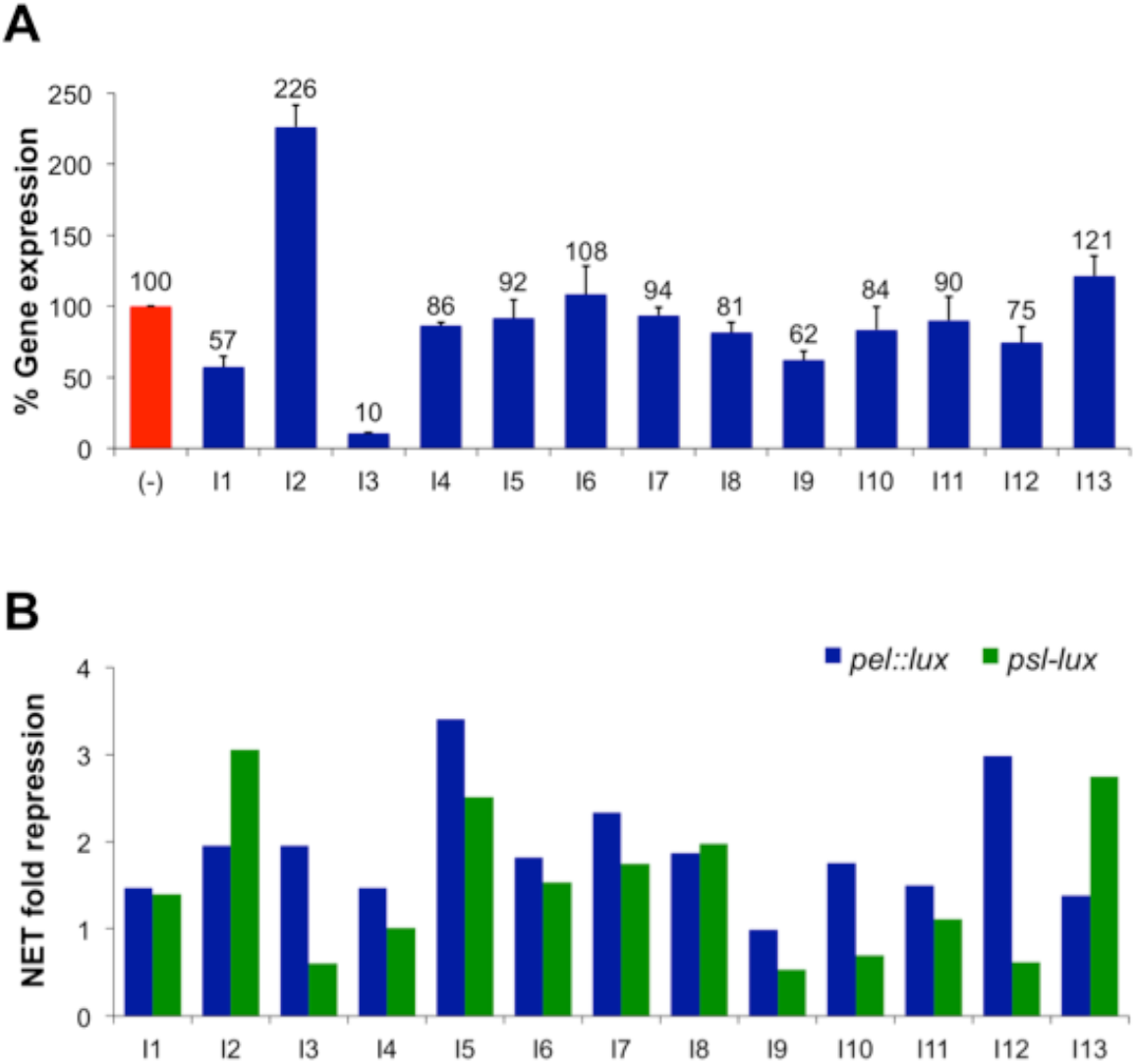
(A) Analysis of compound-treated PAO1::p16*Slux* reporter to assess nonspecific effects on *lux* expression. Gene expression after treatment (blue bars) was compared to untreated control (red) and was represented as the total gene expression throughout 18 hours, which was calculated as area under the curve. Values shown are the mean of triplicate samples plus standard deviation, n=6. **(B)** Repression of *pel* and *psl* after normalization to the expression of 16S genes after treatment. Bars represent the average net fold repression on triplicate samples of the target genes relative to the untreated condition, n=2.

**Figure S2:**
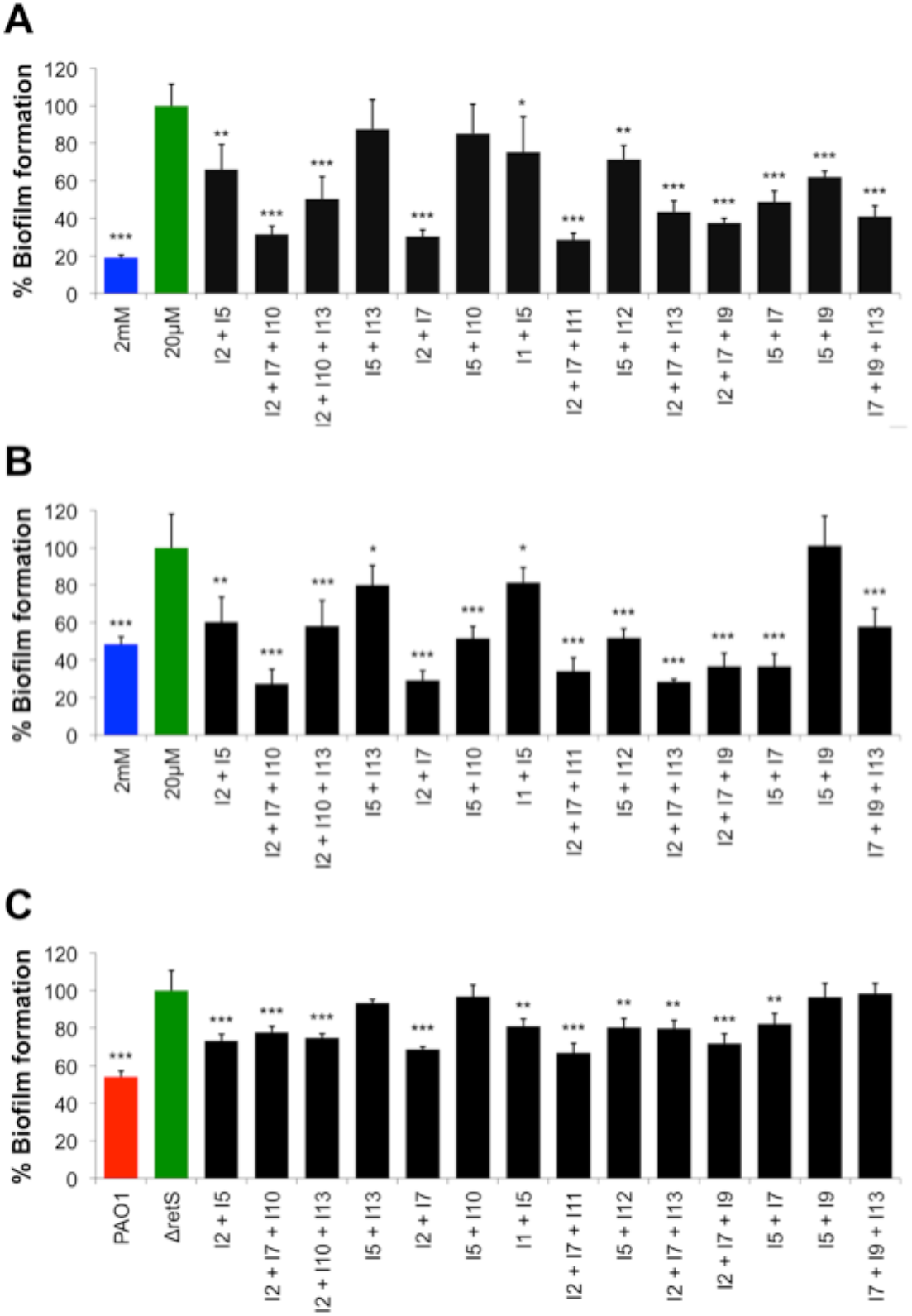
Biofilm assays after combination treatment with select mixtures of antibiofilm compounds. Assays were performed with (A) PA14, (B) PAO1 and the (C) *retS::lux* mutant. Black bars of compound mixture-treated biofilms were compared to green bars of strains grown in biofilm-inducing condition alone. Blue (PA14 and PAO1) and red (PAO1) bars are negative controls of biofilms formed in repressing conditions. Values shown are the mean of 6 replicates plus standard deviation. Significant repression in biofilm formation is indicated: *(p<0.05), **(p<0.01) and ***(p<0.001).

**Figure S3:**
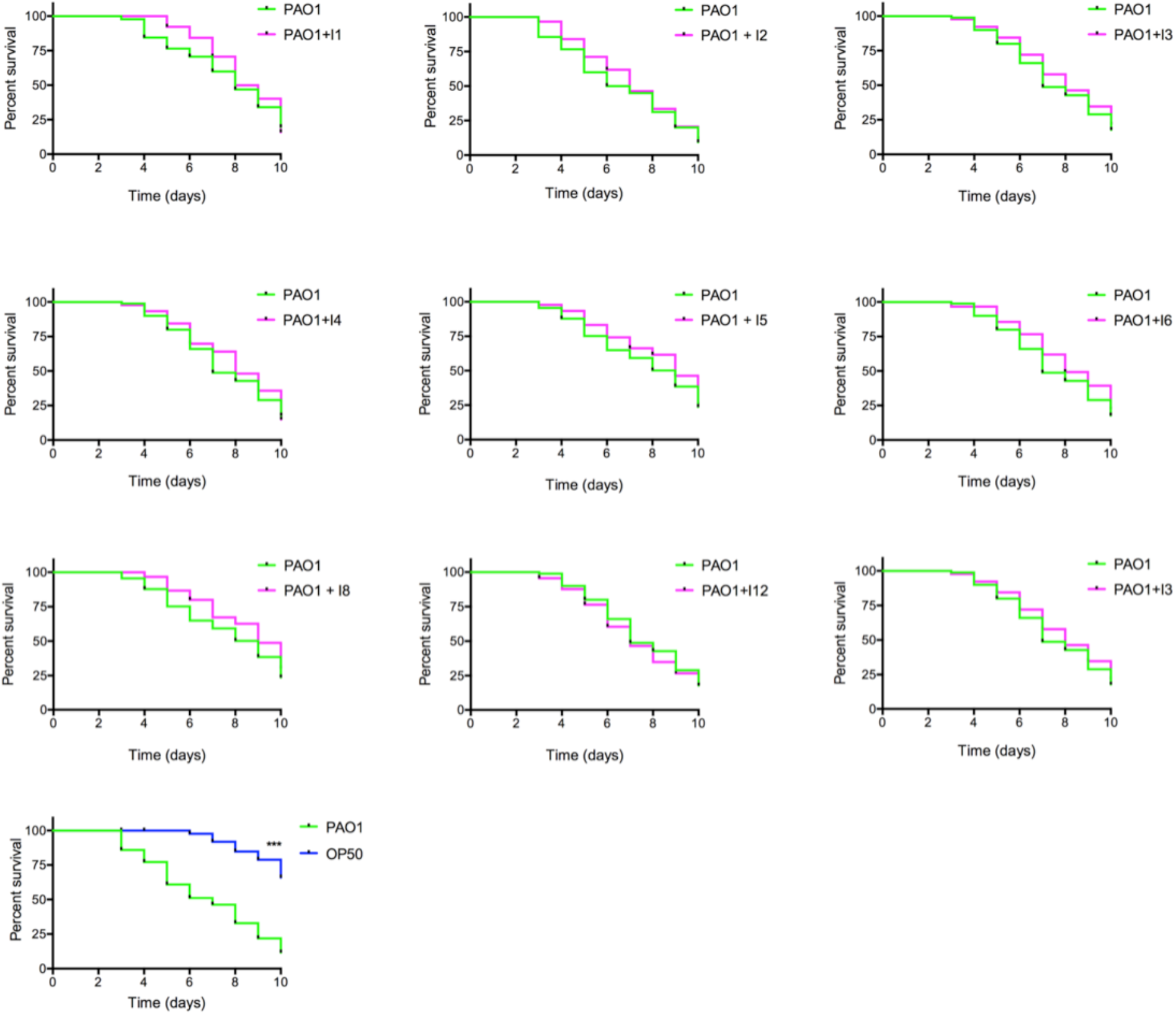
Effects of antibiofilm compounds on the virulence of PAO1 in the *C. elegans* slow killing virulence assay. Kaplan-Meyer curves with % survival represent three independent experiments (n=30) where the total number of worms equals 90. Significance differences in nematode survival are indicated: **(p<0.01) and ***(p<0.001).

**Supplementary Table S1:**
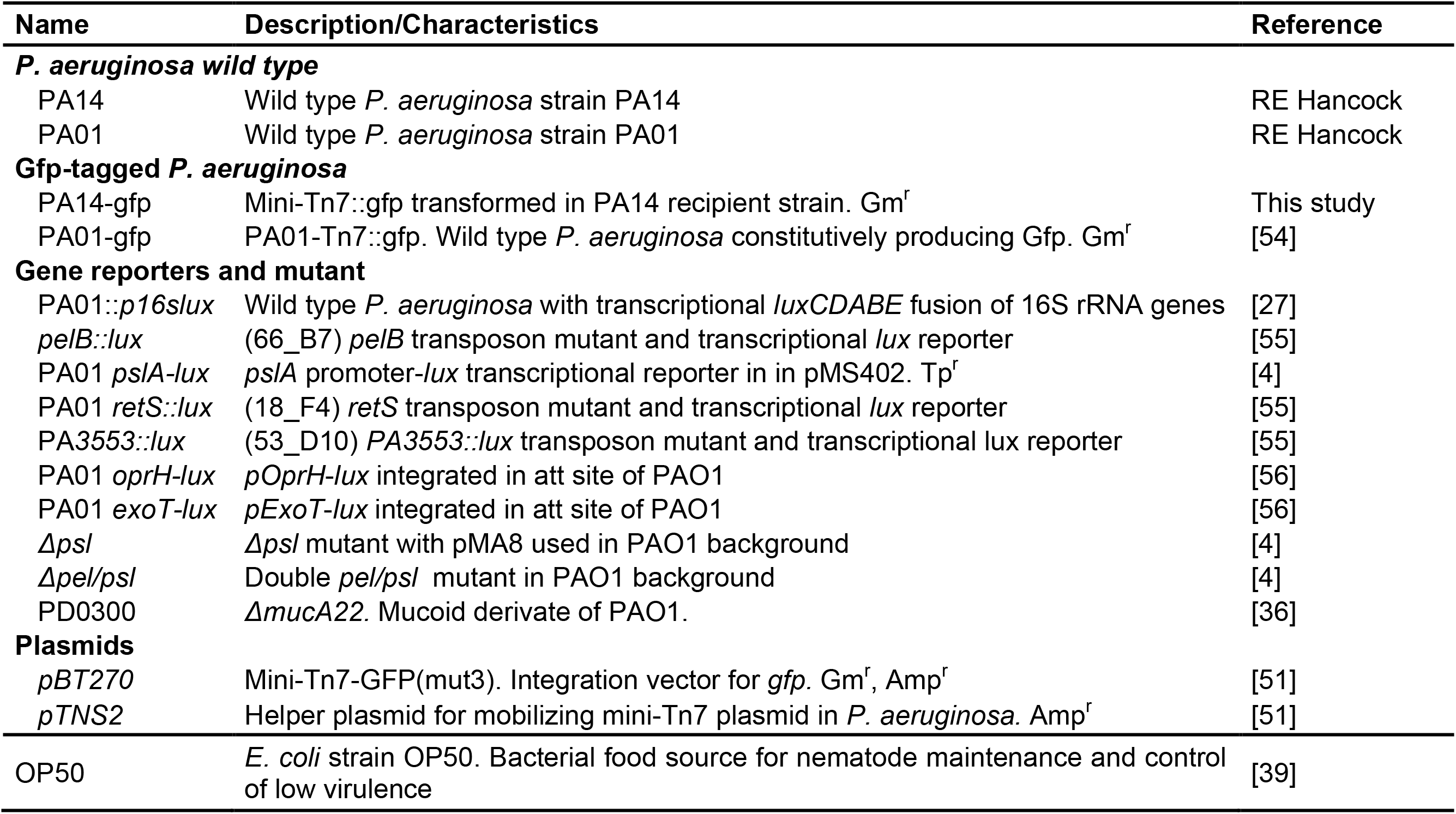
strains and plasmids used in this study.

**Supplementary Table S2:**
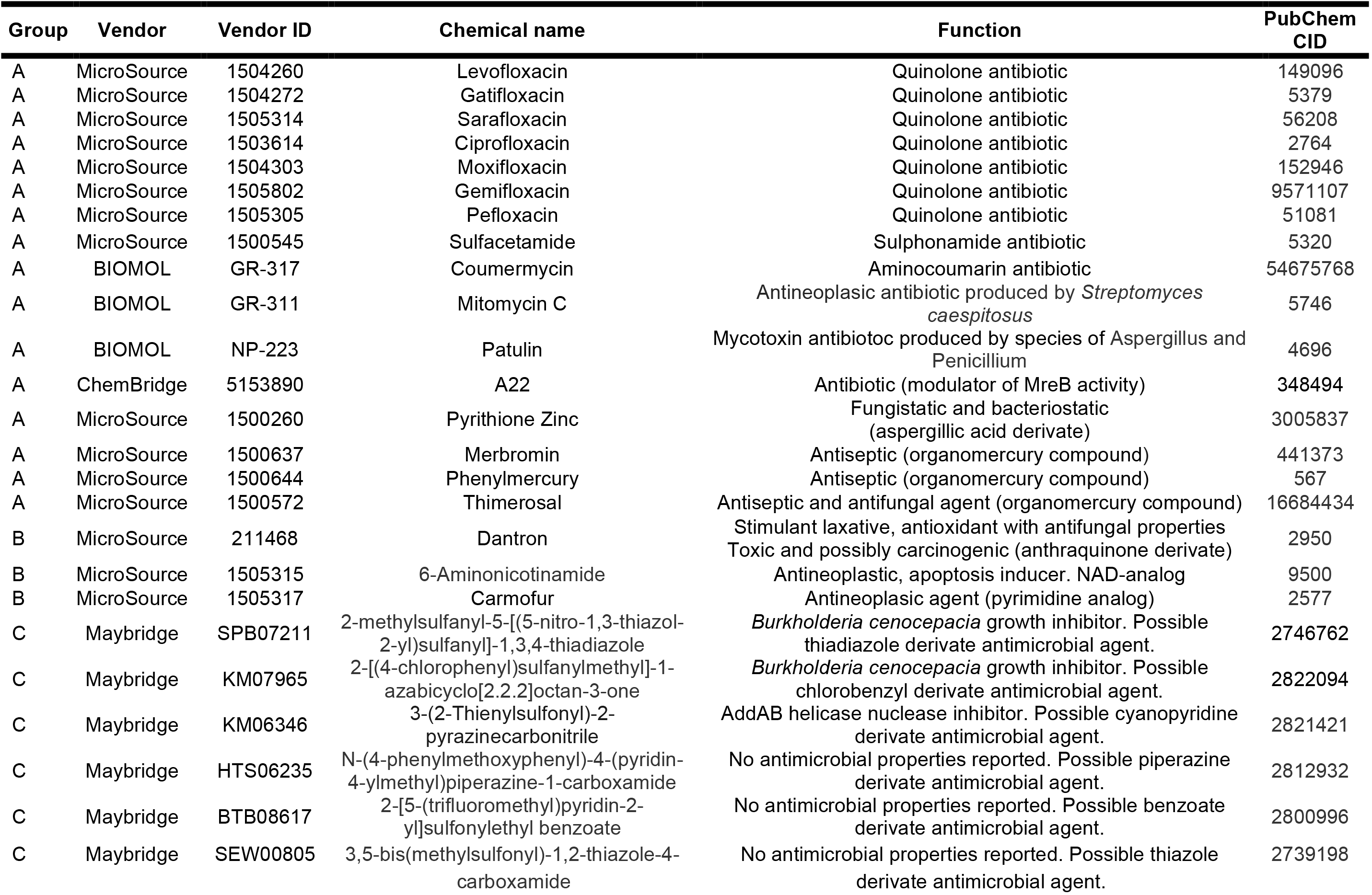
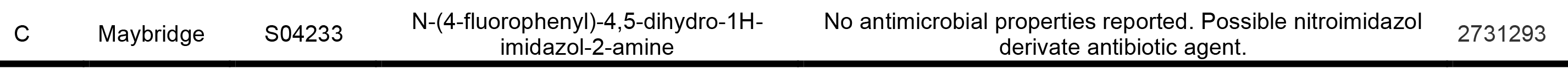
Antimicrobial compounds identified in the HTS screen for EPS repressors.

